# Predictions of bimanual self-touch determine the temporal tuning of somatosensory perception

**DOI:** 10.1101/2024.06.25.600596

**Authors:** Noa Cemeljic, Xavier Job, Konstantina Kilteni

**Affiliations:** Department of Neuroscience, Karolinska Institutet, Stockholm, Sweden; Donders Institute for Brain, Cognition and Behaviour, Radboud University, Nijmegen, The Netherlands

## Abstract

We effortlessly distinguish between touching ourselves with our hands and being touched by other people or objects. Motor control theories posit that this distinction is made possible by the brain predicting the somatosensory consequences of our voluntary movements based on an ‘efference copy’, and attenuating our responses to the predicted self-touch. However, it remains unclear how these predictions impact somatosensory perception at times other than during self-touch: for example, as our hand reaches to touch our body or moves away from it. Here participants discriminated forces applied on their left index finger by a motor. The forces were applied during the reaching movement of their right hand towards the left hand, including the time the reaching ended by simulating self-touch between the hands, or after the reaching movement. We observed that the forces on the left hand felt progressively weaker during the reaching phase, reached their minimum perceived intensity at the time of self-touch, and quickly recovered after the end of the reaching. All effects were replicated with a new cohort of participants that further demonstrated that this gradual attenuation of the perceived magnitude of touch vanished during similar right hand reaching movements that did not produce expectations for self-touch between the two hands. Together, our results indicate a temporal tuning of somatosensory perception during movements to self-touch and underscore the role of sensorimotor context in forming predictions that attenuate the intensity of self-generated touch.

## Introduction

Self-touch occurs in a wide range of everyday activities, from rubbing our eyes with our fists to alleviate itching, to warming ourselves with our hands during shivering, and affectively stimulating our bodies for pleasure. Self-touch behaviors begin in utero ^1,2^ and continue after birth as a way for infants to explore their bodies ^3–5^. Whether spontaneous (*e.g.*, clenching our fists when nervous) or purposeful (*e.g*., moving the hair away from our face) ^6^, self-touch is considered fundamental for building and maintaining a sense of self ^7–10^ through detecting and maintaining contingencies between our motor (*i.e.*, moving to touch our body) and somatosensory cues (*i.e.*, receiving the corresponding touch).

Besides self-touch, we also experience numerous instances of touch generated by other people or objects. Remarkably, our brain possesses the capacity to distinguish between self-touch and touch of external origin to guide behaviour; for example, we do not react to self-touch between our hands, but we might recoil from similar touch initiated by a stranger, just as we would brush away a fly that lands our hand.

How does the brain implement this distinction? Motor control theories posit that the brain uses internal models to predict the somatosensory consequences of voluntary movements ^11–15^ based on a copy of the motor command (*i.e.,* efference copy) and attenuates (or cancels) the responses to the self-generated touch ^16–21^. Accordingly, when reaching with the (active) right hand to touch the (passive) left hand, the internal model predicts and attenuates the resulting touch on both hands using the efference copy associated with the right hand’s reaching movement. In contrast, a touch of external origin cannot be predicted based on the movement and is not attenuated ^17,20^. It has been proposed that this attenuation of self-generated touch facilitates the distinction from externally generated touch by increasing the salience of the latter relative to the former ^15,16,20^. In agreement with this framework, several studies have shown that self-touch elicits an attenuated perceptual response (*i.e.*, it feels less intense or ticklish) ^22–39^ and triggers weaker activity in somatosensory cortices and the cerebellum ^40–46^ than identical touch applied by another person or a machine.

Nevertheless, it remains unclear how the predictions of the internal model about the somatosensory consequences of the movement affect perception at times other than the exact moment of self-touch. For example, imagine reaching with the right hand to touch your left hand (**Figure 1a**). It remains elusive how touch on the left hand is perceived as the reaching movement of the right hand progresses, before contact between the hands occurs, and after the reaching movement has ended. This is due to several reasons. First, several earlier studies assessed somatosensory perception across time on the active hand, rather than the passive hand which served as a target of the active hand’s movement (*e.g.*, ^47–50^). However, it has been demonstrated that somatosensory perception on a limb that moves can be ‘gated’ by nonpredictive mechanisms ^51–57^ (see ^35^ for discussion). To isolate the effects of the internal model’s predictions on somatosensation, other studies have focused on the perception of the passive hand, which is not influenced by non-predictive confounds because it does not move ^16,17,35^. Nevertheless, these studies either probed perception only *at the moment* of self-touch ^22–24,28–31,33–39,58^, overlooking the periods before or after the hands’ contact, or otherwise focused on the time *before* the contact ^59^, overlooking the periods during or after self-touch. Third, other studies have introduced temporal delays to test somatosensory perception before and/or after the predicted self-touch ^26,27,32,46,60^, but those probed timings were tailored to the moment of the predicted tactile sensation and did not directly relate to the reaching movement of the active hand. For example, one can probe somatosensory perception at a fixed time before the anticipated self-touch (*e.g.*, −300 ms), but that timing could align with the early (*i.e.*, acceleration) phase of a short reaching movement (*e.g*., 500 ms) or the late (*i.e.*, deceleration) phase of a longer reaching movement (*e.g.*, 1500 ms). Related to this point, an earlier study ^59^ suggested that somatosensory perception on the passive hand changes only when the active hand reaches its peak velocity. Finally, in addition to being constrained in scope, the abovementioned research has produced conflicting and inconclusive results. Specifically, an earlier study reported a constant increase in the attenuation of touch on the passive hand as the active hand approached the passive ^27^, while another study found facilitation, rather than attenuation, of tactile perception on the passive hand when the active hand reached its peak velocity ^59^.

**Figure 1.**
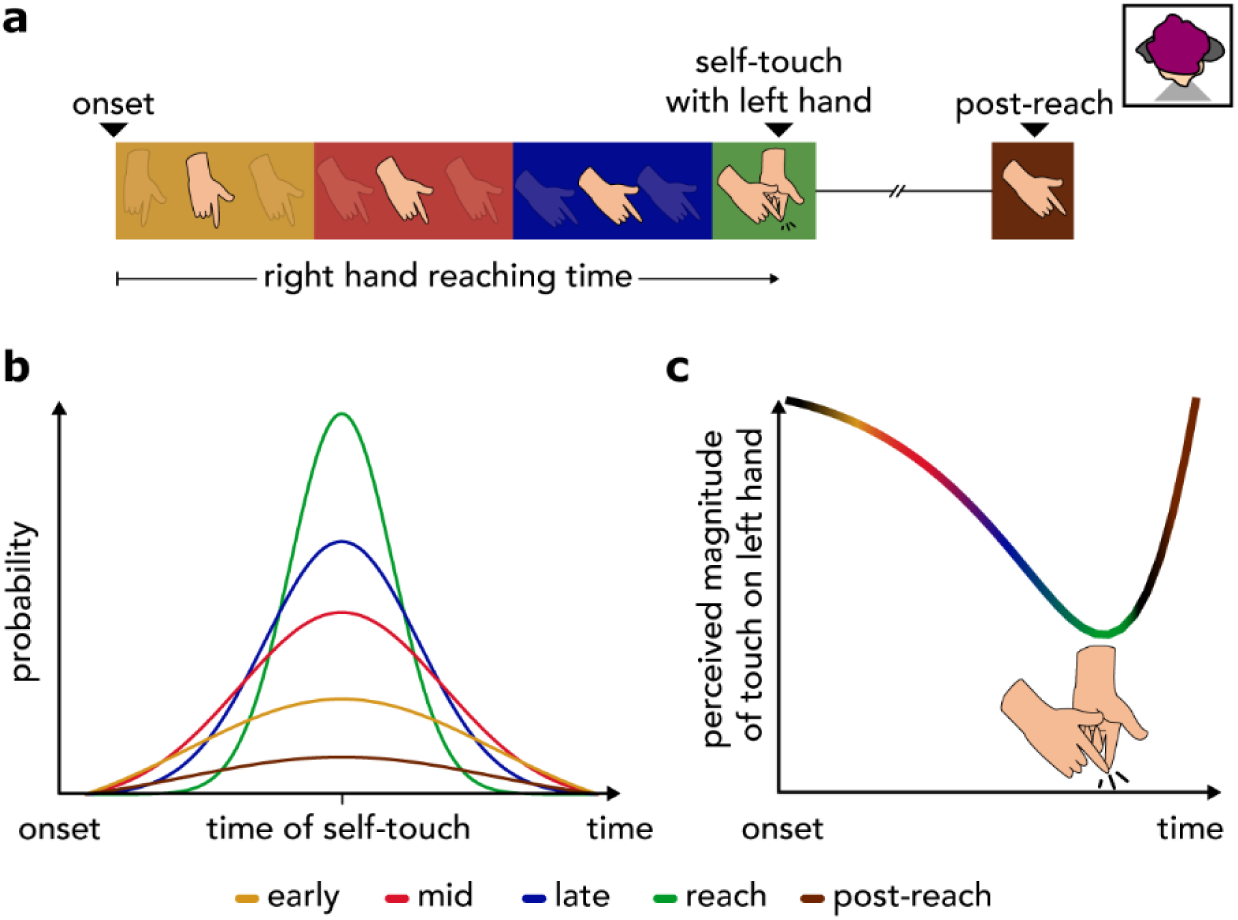
Schematic illustrations of the experimental hypothesis. **(a)** We investigated how participants perceive touch on their left hand during and after reaching with their right hand to touch their left hand. The reaching movement is shown from a third person perspective and the top right small window indicates the orientation of the participant. (**b**) From a theoretical perspective, during the reaching the brain estimates the state of the moving hand (*e.g.*, its position, velocity) using the predicted and received sensory input and updates its predictions about the rest of the movement accordingly. Consequently, as the action progresses, the predictions of the internal model about the timing of self-touch become narrower given that there is a better estimation based on the latest sensory input (*e.g.*, visual and proprioceptive information). (**c)** Touch received during the reaching (yellow, red, and blue) including its conclusion (green), or after the reaching (brown) would be attenuated proportionally to its probability of occurring at that time point. (**a, b, c**) Phases of the reaching movement and times are colour-coded. Yellow represents the early reaching phase (acceleration), red indicates the phase around the peak velocity, blue denotes the late reaching phase (deceleration), green corresponds to the moment the reaching movement concludes upon reaching the target (self-touch), and brown refers to the phase after the reaching.

Here, we hypothesized that the predictions of the internal model should continuously influence somatosensory perception during the reaching movement of the right hand to touch the left hand. Specifically, we reasoned that as the reaching movement of the right hand progresses, the brain gains access to the latest sensory information to adjust the movement and re-estimate the position and velocity of the right hand ^15,19,61^, as well as to update the predictions of the internal model about the remainder of the movement ^62–64^, including the expected timing of contact with the left hand (*i.e.*, self-touch). Consequently, the prediction about the time of self- touch should become sharper (*i.e.* narrower probability distribution) at later timings during the reaching than earlier ones (**Figure 1b**). Therefore, we expected to observe a gradually increasing attenuation of the perceived magnitude of touch during the reaching, which recovers after the hands make contact (**Figure 1c**). That is, the attenuation of somatosensory perception should not be an on-off phenomenon that manifests only at the time of the self-touch, but rather it should gradually build up over the reaching time.

## Results

### Experiment 1. The perceived magnitude of touch gradually decreases during the reaching movement, attaining its minimum at the moment of the predicted self-touch

Twenty-nine participants (15 female, 28 right-handed, aged 18 - 34 years) rested their left hand palm up with their left index finger placed inside a moulded support on a desk. Their right hand was placed palm down on top of a wooden surface placed on top of the desk (**Figure 2a, Supplementary Figure S1**). Upon presentation of an auditory GO cue, they performed a reaching movement finger with their active right hand starting from a designated starting position (reaction time, mean ± SD, 220 ± 42 ms) (**Figure 2a**, *starting position*), and concluded the movement by tapping with their right index finger a force sensor (force duration, mean ± SD, 129 ± 39 ms) placed above, but not in contact with, their left index finger resting on the desk (**Figure 2a**, *cyan force sensor*).

**Figure 2.**
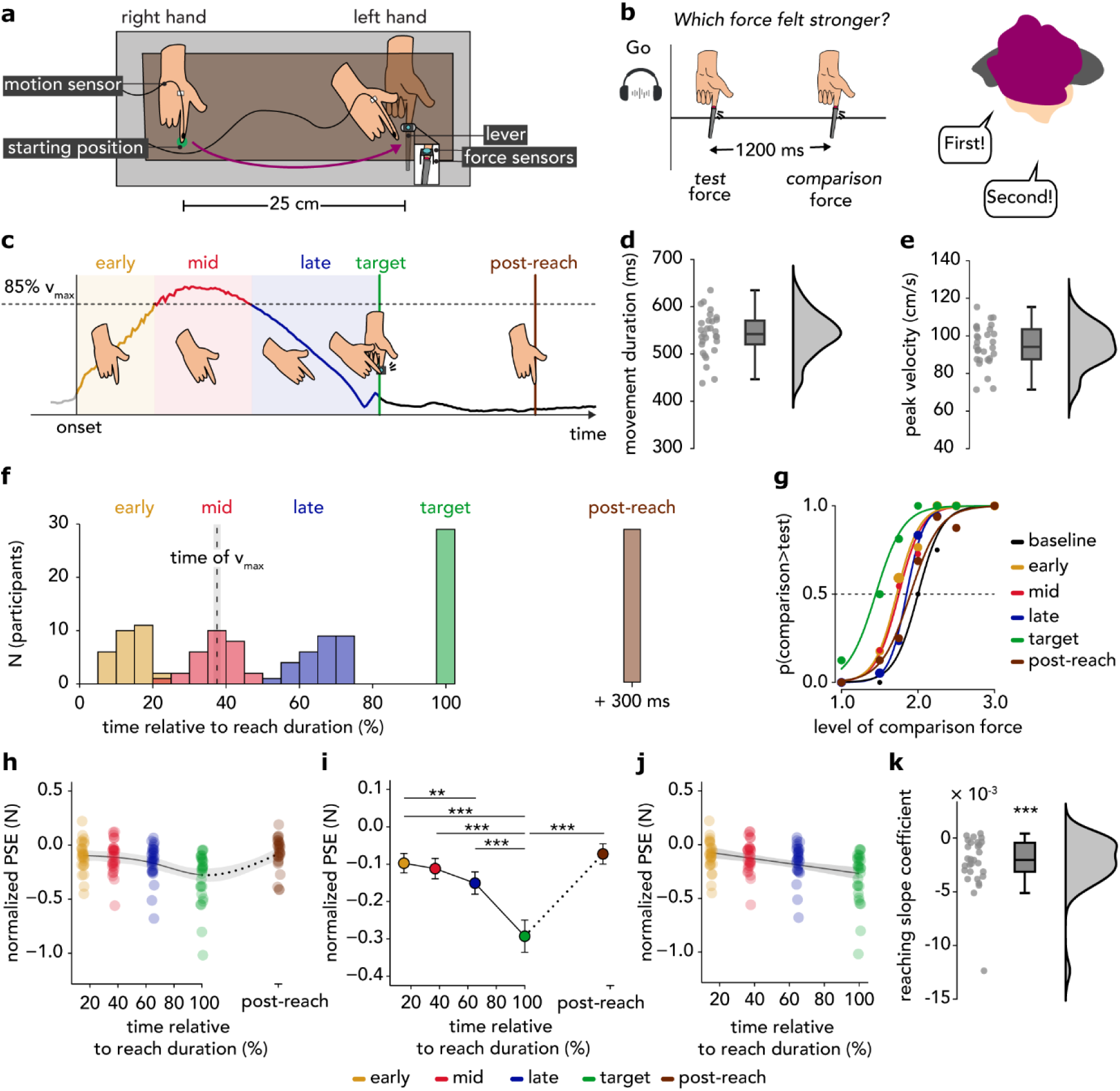
Methods and results of Experiment 1. **(a)** Participants made a reaching movement with their active right hand from the *starting position* to tap the force sensor (depicted as the cyan sensor) placed above but not in contact with their left passive index finger. Another identical force sensor (depicted as the red sensor) was placed within the probe of the lever controlled by the motor to measure the forces applied on the left index finger. In every trial, a motion tracking sensor (*motion sensor*) recorded the position of the right index finger. The right hand was placed on top of the wooden table (brown surface), which was placed on top of the desk (gray surface). For illustration purposes, the wooden table appears transparent. The reaching movement is shown from a third-person perspective. **(b)** In every trial, the participants received two forces on the pulp of their left index finger – the *test* and the *comparison* force – via the lever of the motor. After receiving both forces, they verbally indicated which force felt stronger. **(c)** Example of a velocity profile of one pilot participant illustrating the different phases for a reaching trial. The various phases were determined using the 3D velocity data, but the horizontal velocity is plotted here for simplicity. In the reaching trials, participants received the *test* force on their left index finger during one of five different phases relative to their movement: *early* (yellow), *mid* (red), *late* (blue), at the *target* of the movement (green), or 300 ms *post-reach* (brown). The *comparison* force was always delivered 1200 ms after the *test* force and when the right hand had completed the reaching and had stopped moving. In the *baseline* trials, participants kept both hands relaxed and received the *test* and the *comparison* forces at rest. **(d-e)** Individual and group reach durations **(d)** and peak velocities **(e)**. The markers represent individual data, boxplots show the medians and interquartile ranges, and raincloud plots depict the data distributions. **(f)** Histograms of the times at which the *test* force was applied to the participants’ left index finger in the different types of reaching trials. Times are expressed as percentages relative to the reaching movement duration (*i.e.*, the time needed to tap the force sensor) and are binned into 5% intervals. The vertical dashed line represents the mean time of peak velocity (v_max_), and the grey shaded area represents the standard error of the mean (s.e.m.). **(g)** Responses and fitted logistic models for all types of trials for one participant. We extracted the PSE (*p* = 0.5) for each participant and each trial type. **(h)** Individual PSE values are shown for each type of reaching trial after being normalized to the *baseline* PSE. The individual PSEs of the *early*, *mid*, *late*, and *target* trials are plotted against the time the *test* force occurred in each trial type relative to the reach duration (%). In the *post-reach* trials, the PSEs were computed for the forces delivered 300 ms after the end of the reach (the dotted line indicates a discontinuity in the x-axis). In the figure, individual points are jittered (± 1%) to avoid complete overlap. To illustrate the pattern in our data and facilitate the comparison with our hypothesis (Figure 1c), we overlaid a regression line based on a generalized additive model. The shaded area represents a 95% confidence interval. **(i)** Normalized group PSE values for each trial type (mean ± s.e.m.). The touch felt significantly weaker as the reaching time progressed (*late < early*, *target < early*, *target < mid*, *target < late*) but quickly recovered afterwards (*post-reach > target*). Asterisks denote significant pairwise comparisons of interest after corrections for multiple comparisons. The dotted line indicates the discontinuity in the x-axis. **(j)** Normalized individual PSE data during reaching (*early* to *target*). Data are jittered (± 1%) to avoid complete overlap. The PSEs of each participant were fitted with a linear model, and we display the group’s linear fit. The shaded area depicts a 95% confidence interval. **(k)** Slope coefficients of the participants’ linear models showing the change in PSEs over the reaching. The slopes were overall negative, indicating an increasing attenuation, and significantly different from zero. The markers represent the individual slope coefficients, boxplots show the medians and interquartile ranges, and raincloud plots depict the coefficients’ distributions (** *p* < 0.01, *** *p* < 0.001).

During or after the reaching movement, the participants received a brief force on their left index finger via a probe attached to the lever of a DC electric motor (**Figure 2a**, *lever*). This *test* force had a fixed amplitude of 2 N. We quantified how strongly participants perceived the intensity of this *test* force over time using a well-established force discrimination task ^16,27–35,46^. In each trial, participants received the *test* force (2 N) and a *comparison* force with an intensity that pseudorandomly varied around the intensity of the *test* force (between 1 and 3 N) (**Figure 2b**). The magnitudes of the *test* and *comparison* forces were confirmed by a second identical force sensor placed inside the probe (**Figure 2a**, *red force sensor*). As mentioned above, the *test* force was applied at a different time relative to the reaching movement of the right hand on any given trial. However, the *comparison* force was always applied 1200 ms after the *test* force – that is, when the right hand had finished the reaching and stopped moving. In other words, our experimental manipulation exclusively involved the *test* force, while the *comparison* force served as a reference. After the presentation of the two forces, participants verbally indicated which one of the *test* or the *comparison* force felt stronger (**Figure 2b**).

The participants performed a block of 560 reaching trials in which the *test* force was delivered during one of five different phases relative to the movement of their right hand in randomized order (**Figure 2c**): (a) the *early* reaching trials included the period between movement onset until the time the velocity reached 85% of its peak value (early acceleration phase); (b) the *mid* reaching trials included the period after the velocity reached 85% of its peak value until the time it dropped below 85% of its peak value (peak velocity phase); (c) the *late* reaching trials included the period after the velocity dropped below 85% of its peak value until before the moment the right index finger tapped the force sensor (late deceleration phase); (d) the *target* trials included the moment when the right index finger tapped the sensor and triggered the touch on the left hand, thereby simulating bimanual self-touch (end of reaching); and (e) the *post-reach* trials included the moment at 300 ms after the tap on the sensor and thus after the reaching movement. Participants were aware that the *test* force could come at different times but did not know when they would receive it on every trial. These 5 trial groups were defined to correspond to different phases of the reaching movement ^65,66^, and capture the time of peak velocity, where changes in the somatosensory perception were previously reported ^59^ and the post-reach phase where changes are typically minimal ^27,32^. On average, participants took (mean ± SD) 542 ± 46 ms to reach their left hand (within-subjects SD of movement time, 77 ± 20 ms) (**Figure 2d**) with a peak velocity of 95 ± 11 cm/s at 203 ± 31 ms after movement onset (**Figure 2e**). The average times at which the *test* force was delivered in the *early*, *mid*, and *late* trials were 81 ± 17 ms, 201 ± 31 ms, and 350 ± 27 ms relative to movement onset, corresponding to 15%, 37%, and 65% of the average reaching time in the *early*, *mid*, and *late* trials, respectively (**Figure 2f**) (**Materials and Methods**). There were no significant differences in the forces participants tapped on the sensor with their right index finger between any trial type (*F*(4, 112) = 1.001, *p* = 0.410, *η_p_*^2^ = 0.035)), confirming that the participants behaved comparably across all reaching trials (**Supplementary Figure S2**).

The participants performed an additional block of 56 *baseline* trials in which the participants remained still while performing the force discrimination task. These trials were conducted either before or after the block of reaching trials in a fully counterbalanced order and served to quantify the participants’ perceived magnitude of the *test* force in the absence of action. For all types of trials (*early*, *mid*, *late*, *target*, *post-reach*, and *baseline*), we fitted the participants’ responses from the force-discrimination task with logistic models (see **Supplementary Figure S3** for all individual fits). All fits were very good (McFadden R^2^ > 0.41). Next, we extracted the point of subjective equality (PSE), which indicates the intensity at which the *test* force felt as strong as the *comparison* force (*p* = 0.5) and represents the perceived intensity of the *test* force per trial type (**Figure 2g**). A lower PSE in one trial type compared to another indicates that the *test* force felt weaker in the former than in the latter trial type.

PSE values of the *reaching* trials were first normalized to the PSEs of the *baseline* trials (PSE_reaching_ – PSE_baseline_) and then subjected to a one-way repeated measures ANOVA with the type of trial (*early*, *mid*, *late*, *target*, *post-reach*) as the within-subjects factor. The type of trial was significant: *F*(2.09, 58.44) = 15.856, *p* < 0.001, *η_p_^2^* = 0.362. Planned pairwise comparisons revealed that the perceived intensity of the *test* force gradually decreased the closer the touch was delivered to the moment of the predicted self-touch (**Figure 2h-i**): specifically, the PSEs were significantly lower in the *late* compared to the *early* trials (*n* = 29, *t*(28) = −2.956, *p* = 0.009 *FDR-corrected*, *CI*^95^= [−0.090, −0.016], *d* = −0.549), in the *target* compared to the *early* trials (*n* = 29, *W* = 11, *p* < 0.001 *FDR-corrected*, *CI*^95^= [−0.234, −0.115], *rrb* = −0.949), in the *target* compared to the *mid* trials (*n* = 29, *W* = 31, *p* < 0.001 *FDR-corrected*, *CI*^95^= [−0.225, −0.097], *rrb* = −0.857) and in the *target* compared to the *late* trials (*n* = 29, *W =* 41, *p* < 0.001 *FDR-corrected*, *CI*^95^= [−0.180, −0.064], *rrb* = −0.811). Differences in the PSEs between *early* and *mid* trials (*n* = 29, *t*(28) = 0.829, *p* = 0.414 *FDR-corrected*, *CI*^95^= [−0.021, 0.050], *d* = 0.154) and between *mid* and *late* trials (*n* = 29, *t*(28) = 1.813, *p* = 0.094 *FDR-corrected*, *CI*^95^= [−0.005, 0.083], *d* = 0.337) were not significant. The lowest PSEs thus occurred at the time of contact between the two index fingers (*i.e.*, *target* trials) when the *test* force was perceived to be significantly weaker than the *test* force at all other probed times. These PSEs quickly recovered in the *post-reach* trials and were significantly greater than in the *target* trials (*n* = 29, *W =* 412, *p* < 0.001 *FDR-corrected*, *CI*^95^= [0.135, 0.268], *rrb* = 0.894).

To further corroborate the gradual decrease in PSEs during reaching, we next fitted each participant’s normalized PSEs in *early*, *mid*, *late*, and *target* trials with a linear regression model (**Figure 2j)**. As hypothesized, the extracted slope coefficients were negative (β = −2.293×10^-3^ ± 2.494×10^-3^) and significantly different from zero (*n* = 29, *W* = 14, *p* < 0.001, *CI*^95^= [−0.003, −0.001], *rrb* = −0.936) (**Figure 2k**), indicating that the *test* force was perceived weaker the closer it was delivered to the expected time of self-touch. Similar results were obtained when fitting the PSEs from the *early*, *mid*, and *late* reaching trials only.

Taken together, the results of Experiment 1 demonstrated that the touch felt gradually weaker as the reaching movement progressed, attained its minimum perceived intensity at the moment of the predicted self-touch, and quickly recovered after the reaching.

### Experiment 2. The gradual attenuation of touch during reaching depends on whether the reaching is expected to produce self-touch

The results of Experiment 1 can be attributed to a dynamic influence of the internal model’s predictions on somatosensory perception during the reaching movement. To rule out that the observed PSE effects are due to other factors related to the movement or the task *per se* (for example, general anticipation of tactile stimulation on the left index finger, unspecific time effects, divided attention to both hands or feelings of agency over the action), we conducted a control experiment. In Experiment 2, we hypothesized that if the temporal tuning of somatosensory perception observed in Experiment 1 stems from the predictions of the internal model, then performing a reaching movement with the right hand that does not have any tactile consequences for the left hand should not modulate somatosensory perception in time. This would occur because, in such a scenario, the internal model would not predict somatosensory input on the left hand based on the efference copy (**Figure 3a**).

**Figure 3.**
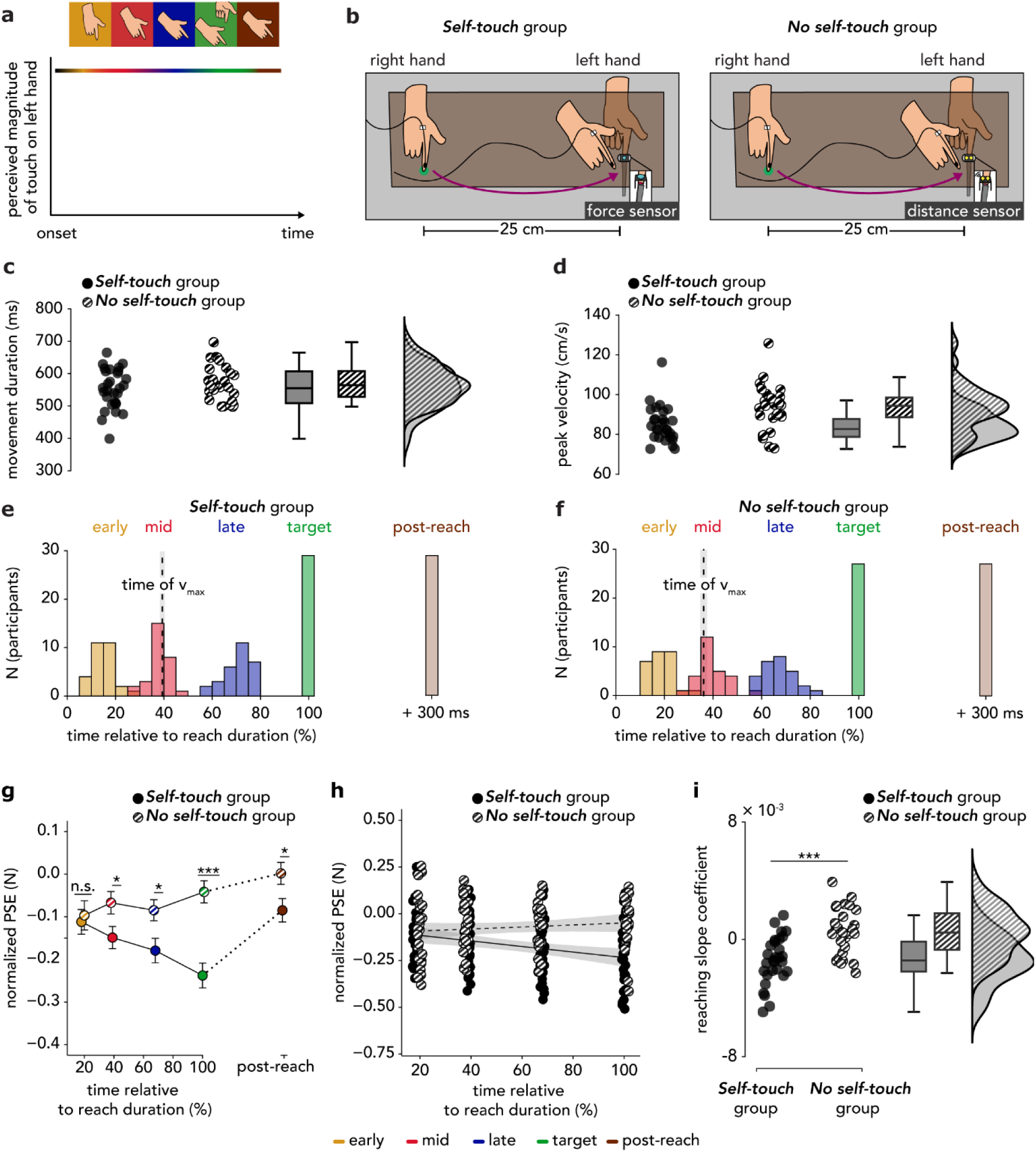
Methods and results of Experiment 2. **(a)** When reaching with the active right hand towards the passive left hand but without planning to touch it (*i.e.*, planning to stop the movement before contact), the brain does not predict somatosensory consequences on the left hand based on the efference copy of the right hand. Consequently, somatosensory perception on the left hand should not show any modulation over time. **(b)** As in Experiment 1, participants in the *Self-touch* group made a reaching movement with their right hand from the *starting position* to tap the force sensor (cyan sensor) placed above, but not in contact with, their left index finger. Participants in the *No self-touch* group (right) also made a reaching movement with their right hand from the *starting position* to approach a distance sensor (yellow sensor) placed above, but not in contact with, their left index finger. The reaching movement is shown from a third-person perspective. **(c-d)** The durations of reaching movements were comparable between the two groups **(c)**. The *Self-touch* group exhibited a lower peak velocity than the *No self-touch* group **(d)**. **(e-f)** Histograms of the times at which the *test* force was applied in the different types of reaching trials for the **(e)** *Self-touch* group and **(f)** *No self-touch* group. The times did not significantly differ between the two groups. Times are expressed as percentages relative to the reaching movement duration and are binned into 5% intervals. The vertical dashed line represents the mean time of peak velocity (v_max_), and the grey shaded area represents the standard error of the mean (s.e.m.). **(g)** PSEs were significantly lower in the *mid*, *late*, *target*, and *post-reach* trials when self-touch was expected (*Self-touch* group) than when it was not predicted (*No Self-touch* group). Asterisks denote between-group comparisons of interest after corrections for multiple comparisons. The dotted line indicates the discontinuity in the x-axis. **(h)** Individual normalized PSE data during the reaching movement only (*early* to *target*) are plotted for each group. Individual points are jittered (± 1%) to avoid complete overlap. We fitted a linear model for every participant in every group separately, but for illustration purposes, we display the group fits. The shaded area depicts a 95% confidence interval. **(i)** In the *Self-touch* group the slopes were overall negative and significantly lower than the slopes in the *No self-touch*, which did not differ from zero. The markers represent the slope value for each participant, boxplots show the medians and interquartile ranges, and raincloud plots depict the data distributions. (* *p* < 0.05, ** *p* < 0.01, *** *p* < 0.001, *n.s.* nonsignificant)

Two new groups of naive participants participated in Experiment 2. Participants in the *Self-touch* group (*n* = 29, 15 female, 28 right-handed, aged 18-37 years) performed the same task as those in Experiment 1: they reached with their right index finger to tap the sensor above their left index finger (reaction time, mean ± SD, 171 ± 38ms) and received forces on their left index finger at different times with respect to the movement of their right hand (**Figure 3b**). In contrast, participants in the *No self-touch* group (*n* = 27, 15 female, 26 right-handed, aged 21- 37 years) performed the same reaching movement with their right hand as participants in the *Self-touch* group did (reaction time, mean ± SD, 130 ± 35 ms), but they were instructed to never make contact between their two fingers through the probe (**Figure 3b**). Rather, they approached a distance sensor placed on top of, but not in contact with, their left index finger which triggered the force once the position of the right index finger became lower than a pre-set position threshold. Then they lifted their right hand as the *Self-touch* group, simulating a virtual tap in the air. The stimuli were delivered at the same time between the two groups, with the *test* force in the *target* trials being applied upon triggering the distance sensor in the *No self-touch* group and upon triggering the force sensor in the *Self-touch* group (see **Materials and Methods**).

As in Experiment 1, participants in the *Self-touch* group were told that their hands would make contact through the probe in some trials, simulating self-touch, but they would not know beforehand which trials these would be. In contrast, participants in the *No self-touch* group were instructed to stop their right hand at a certain distance above their left hand and then lift it without ever making contact between their hands through the probe. Previous studies ^27,28,30^ have shown that if participants deliberately stop their right hand movement above their left hand, the touch applied on the left hand is not attenuated because it cannot be predicted by the motor command. In contrast, if participants predict based on their movement that they will make contact between their two hands but unexpectedly miss the contact, robust attenuation is still observed ^28,30^ indicating that the major driver of somatosensory attenuation is the prediction of self-touch based on the efference copy and not the hands’ contact *per se*. Therefore, in Experiment 2, we expected to replicate all the effects of Experiment 1 with the *Self-touch* group and further observe no temporal modulation in the *No self-touch* group because the internal model would no longer predict touch on the left hand.

The two groups performed reaching movements of comparable durations (**Figure 3c**). The *No-self touch* group exhibited slightly higher peak velocity than the *Self-touch* group (**Figure 3d**), but both groups received the *test* forces at similar times (**Figure 3e-f**, **Supplementary Text S1**). As in Experiment 1, the *Self-touch* group pressed comparable forces across the reaching trials (**Supplementary Figure S4)** and all psychometric fits of both groups were very good (McFadden R^2^ > 0.47) (**Supplementary Figures S5-S6**). Importantly, the *Self-touch* group replicated all effects of Experiment 1 (**Supplementary Text S2, Figure S7**).

To compare the somatosensory perception of the two groups, we inserted the participants’ normalized PSEs in a mixed ANOVA with the group as a between-subjects factor and the type of trial as a within-subjects factor. The model revealed a significant main effect of the trial type (*F*(3.27, 176.33) = 10.044, *p* < 0.001, *η_p_^2^* = 0.157), a significant main effect of group (*F*(1, 54) = 8.284, *p* = 0.006, *η_p_ ^2^* = 0.133) and a significant group-by-trial type interaction (*F*(3.27, 176.33) = 7.072, *p* < 0.001, *η_p_^2^* = 0.116). Pairwise comparisons revealed that the PSEs were significantly lower in the *Self-touch* group than in the *No self-touch* group, in the *mid* (*t*(53.89) = −2.206, *p* = 0.040 *FDR-corrected*, *CI*^95^= [−0.156, −0.007], *d* = −0.590), *late* (*t*(53.37) = −2.488, *p* = 0.040 *FDR-corrected*, *CI*^95^= [−0.171, −0.018], *d* = −0.663), *target* (*W* = 129, *p* < 0.001 *FDR-corrected*, *CI*^95^ = [−0.286, −0.123], *rrb* = −0.670), and *post-reach* trials (*t*(53.97) = −2.299, *p* = 0.040 *FDR-corrected*, *CI*^95^= [−0.162, −0.011], *d* = −0.614). No difference was detected in the *early* trials (*t*(51.84) = −0.338, *p* = 0.737 *FDR-corrected*, *CI*^95^= [−0.105, 0.075], *d* = −0.091) (**Figure 3g)**. In other words, although both groups perceived a comparable magnitude of touch in the early phase of the reaching, their perception deviated at later timings depending on whether the reaching movement had somatosensory consequences for the left hand or not. Further control analyses indicated that the observed PSE differences between the two groups were not due to differences in the kinematics or task demands of the reaching movements (**Supplementary Text S3, Figure S8**).

Next, we fitted linear models to the PSEs during reaching (*early*, *mid*, *late*, and *target* trials) and extracted the slope coefficients for every participant in each group. While the slope coefficients from the *Self-touch* group were negative (β = −1.431×10^-3^ ± 1.622×10^-3^) and significantly different from zero (*n* = 29, *t*(28) = −4.750, *p* < 0.001, *CI*^95^= [−0.002, −8.138×10^-4^], *d* = −0.882), replicating Experiment 1, the slope coefficients from the *No self-touch* group were positive (β = 5.274×10^-4^ ± 1.596×10^-3^) and did not significantly differ from zero (*n* = 27, *t*(26) = 1.717, *p* = 0.098, *CI*^95^= [−1.041×10^-4^, 0.001], *d* = 0.330). As expected, the coefficients were significantly lower in the *Self-touch group* compared to the *No self-touch* group (*t*(53.83) = −4.551, *p* < 0.001, *CI*^95^= [−0.003, −0.001], *d* = −1.217) (**Figure 3h-i**). Similar results were obtained when fitting the PSEs from the *early*, *mid*, and *late* reaching trials only, suggesting that the results are not solely driven by attenuation differences in the *target* trials. Together, the results of Experiment 2 indicate that the gradual attenuation of the perceived magnitude of touch on the left hand depends on whether the reaching movement of the right hand has somatosensory consequences for the left hand (*i.e.*, self-touch).

## Discussion

Determining the time course of somatosensory perception during movements to self-touch is essential for understanding how the brain’s predictions influence perception over time ^15,19,63^. By combining kinematic recordings with somatosensory psychophysics, the present study tested the time course of somatosensory perception during a movement to self-touch; that is, how participants perceived touch on their left hand as their right hand reached to touch the left hand, and moved away from it. Going beyond earlier research that did not test perception during the entire movement of the active hand ^22–24,28–31,33–39,45,58,59^ or tested perception at times that were not linked to the reaching movement of the active hand ^26,27,32,46,67^, our study revealed the timeline of somatosensory perception on the passive left hand during the different phases of the active right hand’s movement to touch it. In Experiment 1, we demonstrated that touch on the left hand was perceived with gradually attenuated intensity as the reaching of the right hand progressed: the touch felt significantly weaker in the late reaching phase (*i.e.,* deceleration) than in the early reaching phase (*i.e.,* acceleration) and was significantly more attenuated at the moment of self-touch. The minimum perceived intensity was observed at the time of self-touch between the hands, and it quickly recovered after the reaching had ended.

All these results were replicated in Experiment 2, where participants in the *Self-touch* group exhibited the same temporal tuning of somatosensory responses. In contrast, participants in the *No self-touch* group who performed similar right hand movements that had no somatosensory consequences for the left hand showed no such modulation. Complementary analysis in that group (**Supplementary Text S2**) showed that the perceived intensity of touch did not change significantly at any probed time during the right hand movement. Together, our findings indicate a temporal tuning of somatosensory perception during movements to self-touch and highlight the role of the sensorimotor context in predicting and attenuating the received sensory input.

Our results are in strong agreement with the computational framework of internal forward models. Specifically, when reaching with the right hand to touch the left hand, the internal model uses the efference copy of the right hand movement to predict the sensory consequences for the right and left hands, including self-touch and its expected timing ^15,17,19^. These predictions are combined with the received sensory input (*e.g.*, visual and proprioceptive) during the movement to re-estimate the state of the right limb *(e.g.*, position, velocity) and update the brain’s predictions about the remainder of the movement ^62–64^. Because both sensory ^18,68,69^ and motor signals ^70–72^ are noisy, the moment of the predicted touch corresponds to a time window reflecting a probability distribution rather than a single point in time and thereby incorporates a certain tolerance for overcoming the intrinsic noise ^17^. As the movement of the right hand progresses, new sensory input is received and the predictions of the internal model become increasingly more precise ^62–64^, narrowing the time window. The received touch is attenuated proportionally to how close they were received with respect to this predicted time window, which continuously narrows during the movement.

Notably, the gradual attenuation of somatosensory perception during movements to self-touch cannot be attributed to task-related differences such as dividing attention between the hands, the mere movement of one hand, a sense of agency, or general cognitive anticipation of tactile stimulation. If this were the case, then we would have observed the same perceptual tuning in the participants of the *No Self-touch* group, in which the same discrimination task and right hand reaching movements were performed. In contrast, we did not observe such a temporal modulation in those participants, and a Bayesian analysis supported the absence of gradual attenuation effects (**Supplementary Text S2**). The direct comparison between the two groups revealed significant differences at almost all probed time points, indicating significantly greater attenuation in the *Self-touch* group than in the *No self-touch* group. This was further supported by linear regression analyses, which showed a significant gradual increase in attenuation during the reaching both in Experiment 1 and in the *Self-touch* group of Experiment 2 but not in the *No Self-touch* group. Finally, further control analyses revealed that these effects cannot be attributed to differences in kinematics or task demands between the two groups.

The results of Experiment 2 demonstrated that the differences in somatosensory perception between the two groups were driven by the sensorimotor context within which participants moved; that is, whether the internal model could predict touch on their left hand based on the motor command of their right hand. Theoretically, the brain should attenuate incoming touch that is likely to be the somatosensory feedback of the movement ^17,19,64^. However, the brain needs to first determine the sensorimotor context that describes the relationship between the movement and the predicted sensory input (*i.e.*, internal model) ^73,74^. This is critical because the same motor command will be associated with different sensory consequences in different contexts. For example, when manipulating objects, the same movement can produce different predictions about sensory feedback in a context where both hands manipulate the same object compared to a context where each hand manipulates a different object ^75^. Similarly, when moving the right hand towards the left hand, the movement of the right hand may predict touch on the left hand in a self-touch context but not in a context where the right hand deliberately stops before contacting the left hand ^27,28,30^. Likewise, pressing the right index finger a few centimetres away from the left index finger should not produce any expectations about receiving touch on the left hand ^17,23,39^ unless in a context where the right hand holds a tool to directly press on the left one ^38^. Consequently, the movements the *No Self-touch* group performed were not associated with any somatosensory consequences on their left hand, which explains why we did not observe an attenuation of the perceived magnitude during their reaching movements. Our results therefore provide evidence that the brain evaluates the sensorimotor context to dynamically shape somatosensory perception in time depending on whether somatosensory input can be predicted based on the efference copy. This conclusion complements changes in movement strategies that humans can make to minimize movement-related suppression effects (*e.g.*, moving with slow velocity ^76^) and attenuation of self-generated input (*e.g.*, applying weak forces that are subject to less attenuation ^77^) when tactile feedback is beneficial for task completion, such as movement guidance ^78,79^ or active object exploration ^80,81^.

Our results cannot be attributed to backward or forward masking of the *test* force by the press of the right index finger. The test force was presented at 192 ms and 174 ms on average for Experiment 1 and Experiment 2 (*Self-touch* group) before the target was reached (*late* trials), and at 300 ms afterwards (*post-reach* trials), while backward masking effects were not detected at 150 ms ^50^ and forward masking effects last only up to 100-200 ms ^82–85^. In addition, our findings do not support the proposal that motor predictions enhance the expected somatosensory signals ^86^, as we did not observe any enhancement effects at any of the time points probed in either of the two experiments. Instead, we consistently observed robust attenuation effects when subjects expected self-touch based on their movement. In line with the present findings, we recently demonstrated that the previously reported “enhancement” effects these proposals are based on ^87^ stem from a biased reference condition used rather than action prediction ^30^. If a bias-free condition is employed, no enhancement effects are experimentally observed ^30^. Moreover, our effects contradict the proposal that attenuation effects are due to non-predictive mechanisms related to double tactile stimulation of both fingers ^87^ as we observed somatosensory perception at times *before* the two hands made contact (*i.e.*, in *mid* and *late* trials), in line with earlier findings ^27^ (see also ^21^). Critically, our effects were dependent on whether participants expected self-touch (*Self-touch* group) and extended beyond the mere influence of the movement of the right hand, the touch on the left hand, or the discrimination task *per se* (*No self-touch* group). Consistent with the current results, we recently demonstrated that attenuation effects are observed independently of the double tactile stimulation of the two hands: touch is robustly attenuated if participants predict that they will make contact between their two hands but unexpectedly miss the contact ^30^ (see also ^28^ for similar findings). Therefore, the present study provides additional evidence that somatosensory attenuation is driven by the prediction of the sensory consequences of voluntary movement rather than somatosensory reafference *per se*.

A methodological strength of the present study is that it assessed how the predictions of the internal model affect the somatosensory perception of a stationary limb. Previous studies have shown that somatosensory perception is ‘gated’ on a limb that moves compared to one at rest^47,50,51,76,88–92^ and that this gating can be variably affected by movement (compare ^59,93–96^).

However, several studies have consistently demonstrated that these gating effects are also observed during passive movements (*i.e.*, in the absence of motor signals), indicating that somatosensory effects on a moving limb are influenced by postdictive mechanisms related to peripheral reafference ^52–55,57,97^. In contrast, somatosensory perception on a limb that remains stationary is not subject to gating effects ^55,92,95,97–99^. Moreover, somatosensory perception on a passive limb is attenuated only if the received touch results from an active movement, not from a passive movement ^34^ or passive stimulation ^27^ (see also ^35^). Thus, our approach isolated the effects of sensorimotor predictions while avoiding the confounding influence of peripheral mechanisms.

Our results indicate a scaling of attenuation magnitude as the reaching movement progresses. In contrast, an earlier study reported no changes in somatosensory perception on the static left hand during the early or late phases of the reaching movement of the right hand, except for tactile facilitation—rather than attenuation—around the time the right hand reached its peak velocity ^59^. We did not observe changes in somatosensory perception depending on when the active hand reached its peak velocity in either of our two experiments, despite having a greater sample size than this earlier study. A reason for this discrepancy could be that we used suprathreshold force stimuli while that study used near-threshold vibrotactile stimuli ^59^. In the auditory domain, studies using near-threshold and suprathreshold stimuli have shown different perceptual effects ^100,101^. However, this remains a speculation as there is no evidence for a similar effect in the tactile modality. Another difference between the two studies is that the earlier study ^59^ was conducted in darkness, while our participants could use vision to guide their movements. The authors of the earlier research suggested that the darkness facilitated the participants’ somatosensory perception to guide their hand movement ^48,49^. However, we believe this interpretation requires further evidence given other findings reporting the opposite effect ^96,102^ or null findings ^47,103^ regarding the interaction between vision and somatosensation during arm movements. Finally, another possible reason for this discrepancy is that participants in the study of ^59^ reached with their right hand to either their left index finger or thumb while receiving stimulation on their little finger. Consequently, their experimental design introduced an incongruency between the finger where touch was expected based on the movement and the finger where touch was received. While the authors argue that tactile perception is uniformly modulated across the whole target hand ^48^ other research has shown that such incongruency can hamper somatosensory attenuation ^104^. Here, in all trials of our experiments, the participants reached with their right hand towards their left index finger while receiving stimulation on their left index finger. Consequently, this allowed us to investigate how somatosensory attenuation on the left index finger changes when the participants form expectations about receiving touch on the same finger.

Understanding how expectations about the somatosensory consequences of the movement (*i.e.*, the internal model) dynamically shape somatosensory perception in time is clinically important, given previously observed alterations in psychosis spectrum disorders ^105–112^, functional movement disorders ^113^, Parkinson’s disease ^36^ and ageing ^37^. Interestingly, patients with schizophrenia show deficits in attenuating self-generated somatosensory input by exhibiting perceptual and neural responses to self-touch that are comparable to those of externally generated touch ^110,111^. Using the same task that was used in the present study, we previously showed that individuals with high schizotypal personality traits, which are considered a subclinical analogue of schizophrenia ^114,115^, also show a reduced ability to attenuate self-touch compared to externally generated touch ^29^. In addition to the somatosensory domain, patients with schizophrenia also show difficulty in attenuating self-generated speech ^116–118^ (see also ^119^) and oculomotor responses ^120^. Together, these findings indicated that individuals with schizophrenia form incorrect predictions about the sensory consequences of their actions, misattributing the received self-generated input to external sources, producing the hallucinations and delusions commonly associated with schizophrenia ^109,112,121–123^. Interestingly, one study showed that patients with schizophrenia have a reduced attenuation of neural responses to self-generated auditory stimuli but a-close-to-normal attenuation for the same stimuli when delivered with a 50 ms delay ^124^, suggesting that the prediction of the sensory consequences of the action might be delayed in patients with schizophrenia ^124,125^ (see also ^126^). Additionally, both patients with schizophrenia ^127–129^ and individuals with high schizotypal traits ^130–133^ tend to exhibit a perturbed temporal profile of multisensory integration, experiencing stimuli that are further apart in time as cooccurring. We therefore speculate that patients may show a different temporal tuning of somatosensory perception during self-touch (*e.g.,* flatter slopes, or shifted temporal window) than that observed in the present study. This hypothesis warrants further exploration in future experiments.

Sensory attenuation is not unique to humans but is observed across many species with research on crickets ^134,135^, mice ^136–138^, weakly electric fish ^139–141^, and primates ^142–144^ showing selective attenuation and even cancellation of neural responses to self-generated sensory inputs. Broadly, this serves to increase the salience and facilitate the interpretation of externally generated sensations ^145–150^. Of particular interest are recent reports of a temporally tuned cancellation of self-produced sounds in mice. Specifically, mice trained to associate tones with moving a lever to a specific position, showed attenuated responses in the auditory cortex to these self-generated sounds when presented at the expected time (*i.e.*, when the lever reached the expected position) ^138^. Importantly, this attenuation was not of fixed magnitude during movement: auditory responses were decreased for sounds played just before movement, or 80 ms after the predicted time or onset and reached their minimum at the expected time ^138^. In addition, mice exhibit attenuated activity in the primary somatosensory cortex when receiving the pulsatile stimulus on their whiskers at the predicted time, while less attenuation is observed if the temporal delays of 50 ms and 100 ms from the predicted time are introduced ^151^. The similarity of these findings with those of the present study suggests the presence of an evolutionarily conserved mechanism for generating precise temporal predictions to attenuate expected sensory inputs. We speculate that such a mechanism could be evolutionarily beneficial by increasing the salience of external stimuli (*e.g.*, enhancing the detection of stimuli an animal receives unexpectedly during movement), thereby improving the detection of potential threats and, consequently, promoting survival.

## Materials and methods

### Participants

Ninety naive participants (47 females and 43 males; 87 right-handed, 2 left-handed, and 1 ambidextrous, aged 18-37 years) were recruited to participate in Experiments 1 and 2. The sample size was set to 30 participants per experiment/group based on our earlier studies investigating somatosensory perception using the same task and equipment ^30–35,46^. After data collection, one participant was excluded from Experiment 1 due to missing kinematic data, and four participants were excluded from Experiment 2: one from the *Self-touch* group because the same response was given in 95% of the *target* trials resulting in an unreliable psychometric fit, and three participants from the *No self-touch* group because they performed very fast right hand movements, resulting in too few *early* trials (< 10 trials, arbitrary threshold) (see *Kinematic trials – segmentation*). Therefore, the analysis of Experiment 1 was based on data from 29 adults, and the analysis of Experiment 2 was based on data from 56 adults: 29 in the *Self-touch* group and 27 in the *No self-touch* group.

In both experiments, handedness was assessed using the Edinburgh Handedness Inventory (mean ± SD, 83 ± 25, range: −60 to 100) ^152^. All participants provided their informed written consent and reported not having any neurological or psychiatric disorders or taking any psychoactive medication. The experiments were approved by the Ethical Review Authority of Stockholm (registration number: 2021-03790).

### Experimental apparatus and procedures in Experiment 1

In both experiments, the participants sat at a desk and rested their left hand palm up, with the left index finger placed inside a support (**Figure 2a**, **Supplementary Figure S1**). A vacuum pillow (AB Germa) was provided to support the participants’ left arm and increase their comfort. On top of the pulp of their left index finger laid a cylindrical probe (20 mm diameter) that was attached to a lever controlled by a DC electric motor (Maxon EC Motor EC 90 flat; manufactured in Switzerland) and operated by an Arduino Mega 2560 microcontroller using custom software written in C++ (**Figure 2a**, *lever*). A force sensor (FSG15N1A, Honeywell Inc.; diameter, 5 mm; minimum resolution, 0.01 N; response time, 1 ms; measurement range, 0–15 N) was placed within the probe to record the forces exerted on the left index finger by the motor (**Figure 2a**, *force sensor* depicted in red). A force of 0.1 N was constantly applied to the left index finger to maintain contact between the probe and the finger. A second identical force sensor was placed inside an aluminium capsule (20 mm diameter) and positioned on top of, but not in contact with, the cylindrical probe (**Figure 2a**, *force sensor* depicted in cyan). The right forearm was comfortably placed on top of sponge padding, and the right hand rested above a wooden table (70 cm × 25 cm × 12.5 cm). The wooden table was placed on the desk in front of the participants covering their left hand, and thus preventing direct visual feedback from the left hand and fingers. The right index finger was placed at a position marked on top of the wooden table (**Figure 2a**, *starting position*), corresponding to 25 cm to the right of the left index finger.

Upon an auditory GO-cue generated by the Arduino, participants were instructed to perform a reaching movement with their right hand from the starting position towards the force sensor located above their left hand. They were instructed to perform a fast but natural movement that would be concluded once they tapped the force sensor with their right index finger (*i.e.*, touching the force sensor with the right index finger and immediately lifting the finger). Participants were allowed to freely look at their right hand and the force sensor as needed to guide their movement. A motion tracking sensor (Polhemus Liberty, Micro Sensor 1.8^TM^) was attached to the right index finger with medical tape to continuously record the right index finger position (**Figure 2a**, *motion sensor*). White noise was provided by a laptop through a pair of headphones to mask any sounds made by the motor.

### Force trials

In each experiment, there were 560 reaching trials and 56 *baseline* trials, resulting in a total of 616 trials per participant grouped in two experimental blocks. The seven possible intensities of the *comparison* force (1, 1.5, 1.75, 2, 2.25, 2.5, 3 N) were tested 16 times for every type of reaching trial and 8 times in the *baseline*. The intensity of the *test* force was always fixed at 2 N and its duration was set to 100 ms.

During the reaching trials, we experimentally manipulated the times at which the *test* force was applied on the left index finger. We applied the *test* force at five different times, based on pilot experiments, to correspond to different phases of a natural right hand movement reaching toward the left hand. The *test* force could be applied at 250 ms, 370 ms, and 550 ms after the go-cue, but the different trials were binned into time bins depending on how the participants moved on every trial (*i.e.*, times of movement onset, peak velocity, movement offset) to account for within- and between- subjects differences in movements. Specifically, based on the time at which the *test* force occurred with respect to the participants’ movement (see *Kinematic trials – segmentation*), the trials were binned to correspond to the *early*, *mid,* and *late* phases of the right hand movement, at the time when the participants tapped the sensor (*target*), which simulated self-touch between the hands (system delay ∼35 ms), and at a fixed time of 300 ms after tapping (*post-reach*). All 5 types of reaching trials were randomly interleaved and had an equal probability of occurrence (20%). During the *baseline* trials, participants remained motionless, and the *test* force was applied 100 ms after the onset of the trial, followed by the *comparison* force 1200 ms later. In every trial, after receiving both forces, participants responded verbally about which of the two forces (the first force - *test* or the second force - *comparison*) felt stronger. Participants were instructed not to balance their responses (*i.e*., saying 50% of the time that the first or the second force is stronger), and were further instructed to make their best guess if the intensity of the two forces felt similar. No feedback was ever provided to the participants regarding their responses.

### Experimental apparatus and procedures in Experiment 2

Experiment 2 was a between-group experiment. The *Self-touch* group (**Figure 3b**) performed the same task as the participants in Experiment 1. For the *No self-touch* group, the force sensor that participants approached with their right index finger (**Figure 3b**, *force sensor* depicted in cyan) was replaced by a distance sensor (Ultrasonic Distance Sensor HC-SR04 5 V Version) (**Figure 3b**, *distance sensor* depicted in yellow). The distance sensor was placed on top of, but not in contact with, the probe and was connected to an Arduino microcontroller that controlled a servo motor (Grove-Servo). Once the distance sensor detected that the position of the right index finger became lower than a pre-set position threshold, it triggered a servo motor that hit the force sensor that triggered the *test* force. The servo motor and the force sensor were hidden from the participants’ view. The position threshold was set to approximately match the position participants had when pressing the force sensor with their right index finger in Experiment 1 and in the *Self-touch* group. A visual marker was placed 2 cm above the distance sensor to indicate the approximate vertical position where participants should perform the virtual tap to avoid physical contact with the sensor, and the position of the pre-set threshold was adjusted to this distance. Participants were instructed to perform a fast but natural movement with their right hand toward the distance sensor and to conclude the movement by tapping the right index finger above the sensor without touching it. That is, the participants were instructed to perform movements like those of Experiment 1, with the difference that they never experienced contact between their two fingers and thus never expected to make such contact. Similar to our previous study ^30^, we demonstrated the movement to the participants prior to the experiment, and there were practice trials to ensure that they performed the movement as requested before starting the experiment. The end of the reaching was defined as the time point at which the distance sensor was triggered by the downward movement of the right index finger.

All probed times in Experiment 2 were identical for the *Self-touch* and *No self-touch* groups. However, forces were delivered 100 ms later in Experiment 2 than in Experiment 1. All the other details, including the analysis, remained identical to those of Experiment 1.

### Kinematic trials – preprocessing

The position of the right index finger was recorded at a sampling rate of 240 Hz. TTL signals were used to indicate the onset of the trial, as well as the *test* force and the *comparison* force, thus co-registering the force trials with the kinematic trials. We smoothed the position data using a moving average filter of 5 data points span (≈ 21 ms).

### Kinematic trials – segmentation

For each reaching trial, we calculated the movement onset as the first time point during the trial when the velocity exceeded 5 cm/s for 100 ms sequentially ^93,95,96^. The end of the reaching (*i.e.*, offset) was calculated as the first time point when the force exerted on the force sensor exceeded a pre-set threshold (0.3 N) indicating that participants tapped the sensor with their right index finger and thus concluded the movement. The duration of the movement was calculated as the time difference between the movement offset and onset.

Next, we calculated the time when the 3D velocity of the right hand reached its peak value. To capture the peak velocity of the reaching movement, which typically occurs during the first half of the movement ^65^, and not the peak velocity of the index finger when tapping the sensor (occurring at the end of the movement), we restricted the time window of the peak velocity to within 67% of the movement duration.

Based on the calculated kinematic parameters of every reaching movement, we binned the reaching trials into three distinct categories depending on when the *test* force occurred within a specific trial: *early* trials (*test* force occurring between the movement onset and the time the right hand velocity reached 85% of its peak value), *mid* trials (*test* force occurring after the right hand velocity reached 85% of its peak value up until it dropped below 85% of its peak value), and *late* trials (*test* force occurring after the right hand velocity dropped below 85% of its peak value and before the right index finger made contact with the force sensor). Therefore, this binning approach allowed us to tailor the probed times to the different phases of the reaching movement, by taking into consideration potential changes in reaction and reaching times from trial to trial. In addition, the trials when the *test* force occurred simultaneously with the participant tapping the force sensor were classified as *target* trials, and the trials when the *test* force occurred 300 ms after the participants’ tap on the force sensor were classified as *post-reach* trials (**Figure 2c**). Therefore, the final five groups of reaching trials entering the statistical analysis were *early*, *mid*, *late*, *target*, and *post-reach* trials.

### Exclusion of trials in Experiment 1

As mentioned earlier, each participant was administered 616 trials resulting in 17864 trials in total (29 × 616). We excluded (a) any *baseline* or reaching trials in which the intensity of the *test* force (2 N) was not properly applied (*test* < 1.85 N or *test* > 2.15 N), or the participants’ force responses or kinematic data were not properly registered, (b) any reaching trials in which the participants failed to tap the sensor or did not press/release the sensor properly, (c) any reaching trials in which the participants did not move as instructed (*e.g.*, they moved before the auditory GO-cue, they started their movement from a position closer to the force sensor than the starting position (< 5 cm away from the starting position on the x-axis)), (d) any reaching trials in which the participants received the *comparison* force while moving, and (e) any *early*, *mid*, or *late* trials in which the *test* force was applied either before the participants started moving (*i.e.*, participants were too slow in starting their movement) or after they tapped the sensor (*i.e.*, participants were too fast in finishing their movement). Finally, since our setup contained metallic surfaces (*e.g.*, the aluminium capsule) that can distort the electromagnetic recordings of the motion tracking sensor when in close proximity, we rejected (f) any reaching trials in which the sensor was inside the distortion range at times used for trial segmentation (see below). In total, we rejected 1590 out of 17864 trials (8.9%). This resulted in an average of 110 *early*, 84 *mid*, 92 *late*, 109 *target*, 110 *post-reach,* and 56 *baseline* trials.

### Exclusion of trials in Experiment 2

We used identical criteria for excluding trials as in Experiment 1 with the exception that in the *No self-touch* group we also rejected any *target* and *post-reach* trials in which the participants did not trigger the distance sensor when moving their finger downwards. In addition, for this group we also modified our exclusion criteria under b) (see above) to exclude any trials in which the distance sensor was not properly triggered, instead of participants failing to press the sensor. In total, we excluded 1411 out of 17864 trials (7.9%) in the *Self-touch* group and 2051 out of 16632 trials (12.3%) in the *No self-touch* group. This resulted in an average of 87 *early*, 101 *mid*, 108 *late*, 107 *target*, 108 *post-reach* trials, and 56 *baseline* trials for the *Self-touch* group and an average of 64 *early*, 114 *mid*, 136 *late*, 85 *target*, 85 *post-reach,* and 56 *baseline* trials for the *No self-touch* group.

### Psychophysical fits

The participants’ responses were fitted with psychometric curves (**Figure 2g**), using the *glm* function of the R *stats* package, with a generalized linear model (Equation 1):

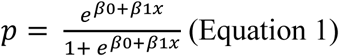

We extracted the point of subjective equality 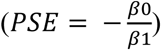 that corresponds to the intensity at which the *test* force felt equally strong to the *comparison* force (*p* = 0.5). This was our variable of interest since it represents the perceived intensity of the *test* force. For control analysis, we also extracted the just noticeable difference in Experiment 2 that reflects the participants’ discrimination capacity in the psychophysics task 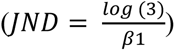 and corresponds to the difference between the thresholds at *p* = 0.5 and *p* = 0.75. A higher JND indicates poorer discrimination ability for the applied forces, suggesting greater difficulty of the task. Before fitting the responses, the *comparison* forces were rebinned to the nearest of seven possible force intensities (1, 1.5, 1.75, 2, 2.25, 2.5, or 3 N). The goodness of fit was assessed using McFadden’s pseudo R^2^.

### Statistical analyses

The data were analysed in R (version 4.2.2) ^153^ and JASP (version 0.17) ^154^. We performed repeated measures and mixed ANOVAs to analyse the data. In the case of sphericity violations, Greenhouse-Geisser corrections were applied. Effect sizes for the ANOVA were given by partial eta-squared (*η_p_^2^*). Planned pairwise comparisons between trial types were performed using parametric (paired or independent-sample *t-*tests) or nonparametric (Wilcoxon signed-rank and Wilcoxon rank sum) tests depending on the normality of the variable distributions. We limited the pairwise comparisons to those central to our hypothesis, which focused on indicating a serial time dependence during reaching (*early* vs. *mid*, *early* vs. *late*, *early* vs. *target*, *mid* vs. *late*, *mid* vs. *target*, *late* vs. *target*) and recovery after reaching (*target* vs. *post-reach*). The normality of the data was assessed with the Shapiro‒Wilk test. Corrections for multiple comparisons were performed using the false discovery rate (*FDR*) and are denoted as “*FDR-corrected*”. For every statistical comparison, we report the corresponding statistic, the 95% confidence intervals (*CI*^95^), and the effect size (Cohen’s *d* or the matched rank-biserial correlation (*rrb*)). Linear regression and multiple linear regression were performed using the *lm* function of the R *stats* package. A Bayesian control analysis was performed to provide evidence in favour of the null hypothesis for comparisons of interest. The analysis was done in JASP using default Cauchy priors of 0.707, where the Bayes factor *BF_0+_* represents the likelihood of the null hypothesis compared to the alternative hypothesis for the one-tailed tests. We interpret a Bayes factor from 1 to 3 as providing ‘anecdotal’ support for the null hypothesis, a Bayes factor from 3 to 10 as providing ‘moderate’ support for the null hypothesis, and a Bayes factor above 10 as providing ‘strong’ support for the null hypothesis ^155–157^. General additive model regression lines were generated using the *geom_smooth* function of the R *ggplot2* package.

## Acknowledgments

The authors would like to thank Dr. Pawel Tacikowski, Dr. Antonella Maselli, Dr. Luc Selen, and Dr. Maximilian Hauser for fruitful discussions. Experiment costs were covered by the Swedish Research Council (VR Starting Grant 2019-01909) and the Åke Wibergs Foundation (M20-0038). N.C. was supported by the Swedish Research Council (VR Starting Grant 2019- 01909). X.J. received funding from the European Union’s Horizon Europe research and innovation program under Marie Sklodowska-Curie (101059348). K.K. was supported by the European Research Council (ERC Starting Grant 101039152).

## Author Contributions

**Noa Cemeljic:** Conceptualization, Data curation, Formal Analysis, Investigation, Software, Visualization, Writing – original draft, Writing – review & editing. **Xavier Job**: Conceptualization, Writing – review & editing. **Konstantina Kilteni:** Conceptualization, Data curation, Formal Analysis, Funding acquisition, Methodology, Software, Visualization, Supervision, Writing – original draft, Writing – review & editing.

