## Supplementary Material for "Predictions of bimanual self-touch determine the temporal tuning of somatosensory perception"

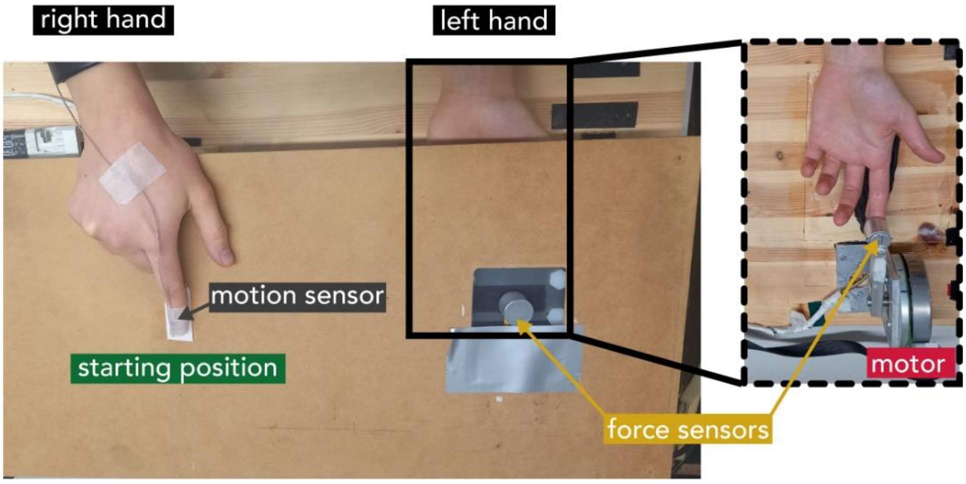


**Figure S1. The experimental setup.** As described in the Methods, the motor, probe, and corresponding force sensor (*right*) were hidden under the wooden table and were not visible to the participants. Participants could only see the sensor they had to tap with their right index finger.

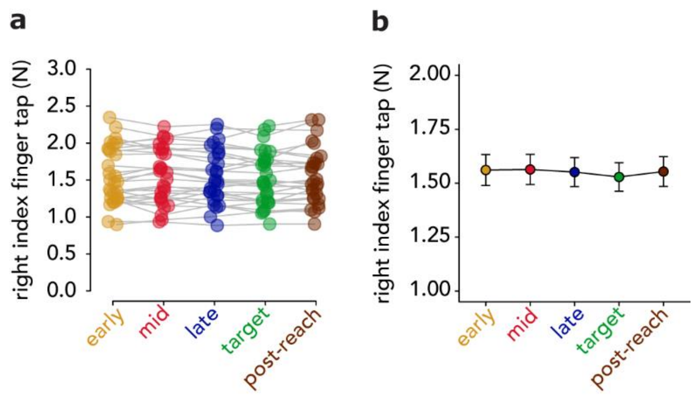


**Figure S2. Forces applied by the right index finger in the reaching trials of Experiment 1.** Individual **(a)** and group **(b)** (mean ± s.e.m.) values for the participants' taps. There was no significant effect of trial type, indicating comparable taps between trials.

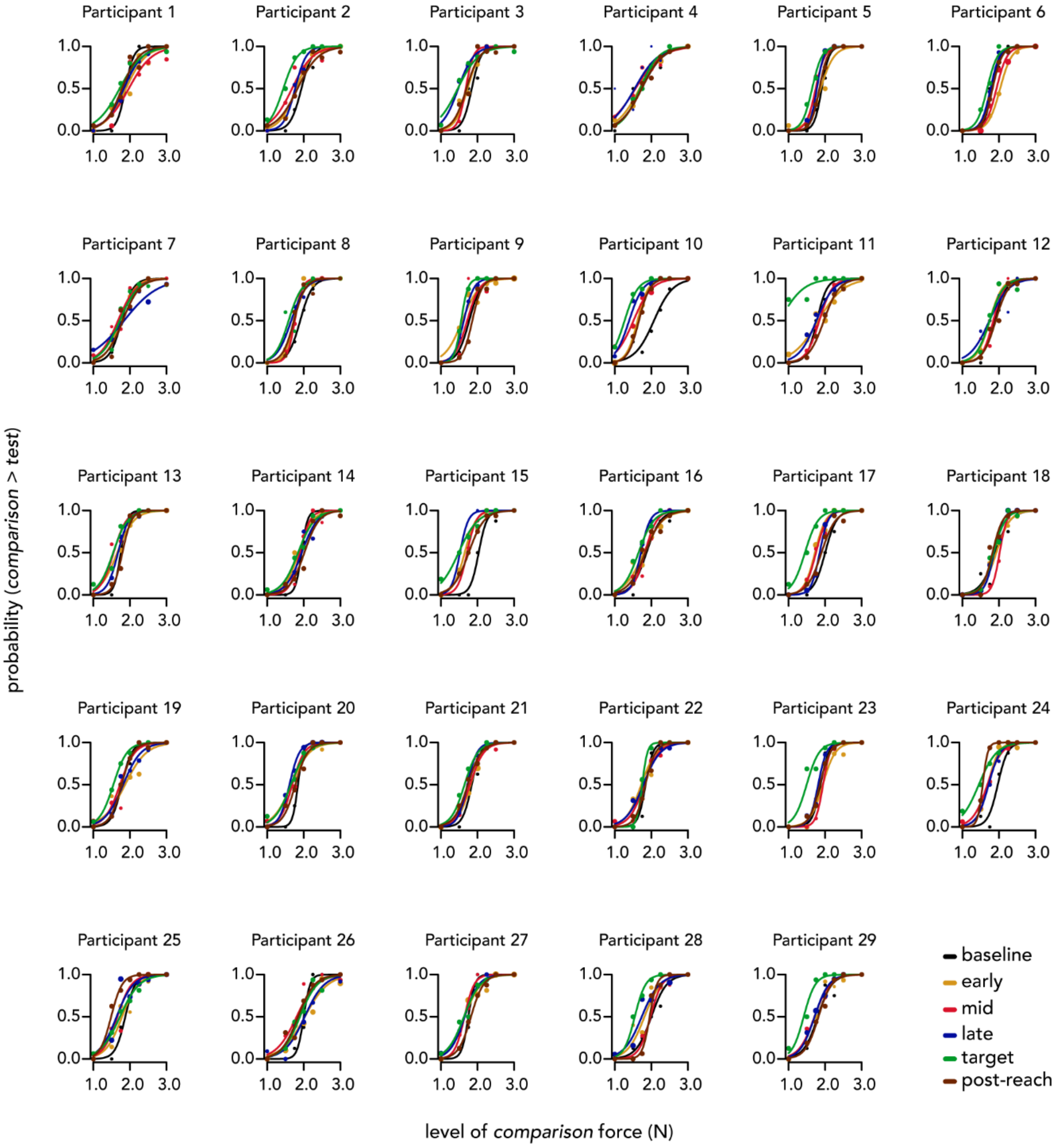


**Figure S3. Fitted logistic models of participants' responses in Experiment 1.**

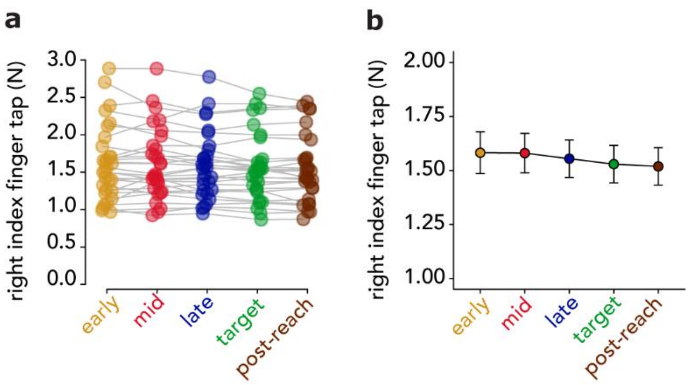


**Figure S4. Forces applied by the right index finger in the reaching trials of Experiment 2 (*Self-touch* group).** Individual **(a)** and group **(b)** (mean ± s.e.m.) values for the participants' taps. There was no significant effect of trial type, indicating comparable taps between the movement trials.

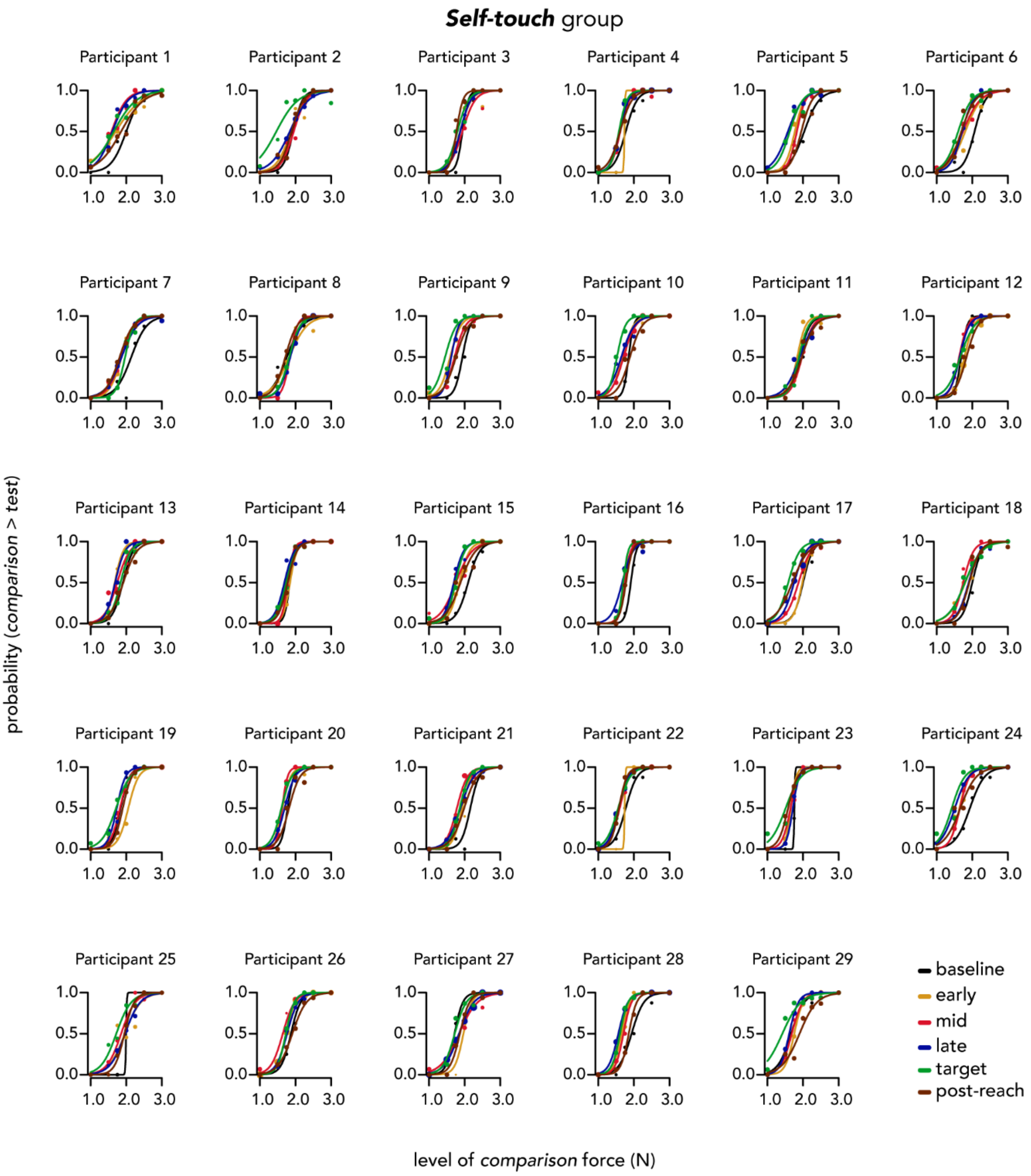


**Figure S5. Fitted logistic models of participants' responses in the *Self-touch* group in Experiment 2.**

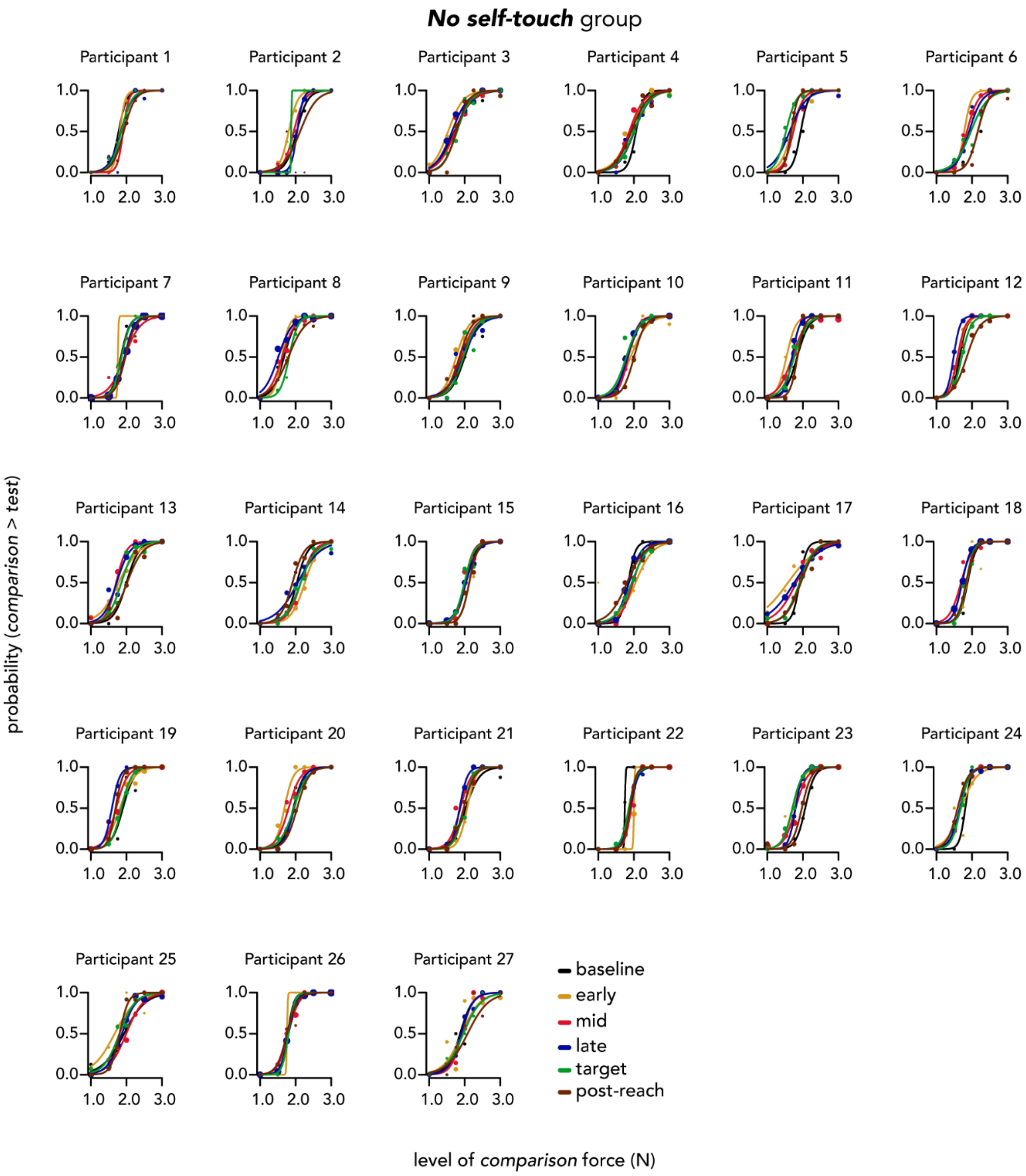


**Figure S6. Fitted logistic models of participants' responses in the *No self-touch* group in Experiment 2.**

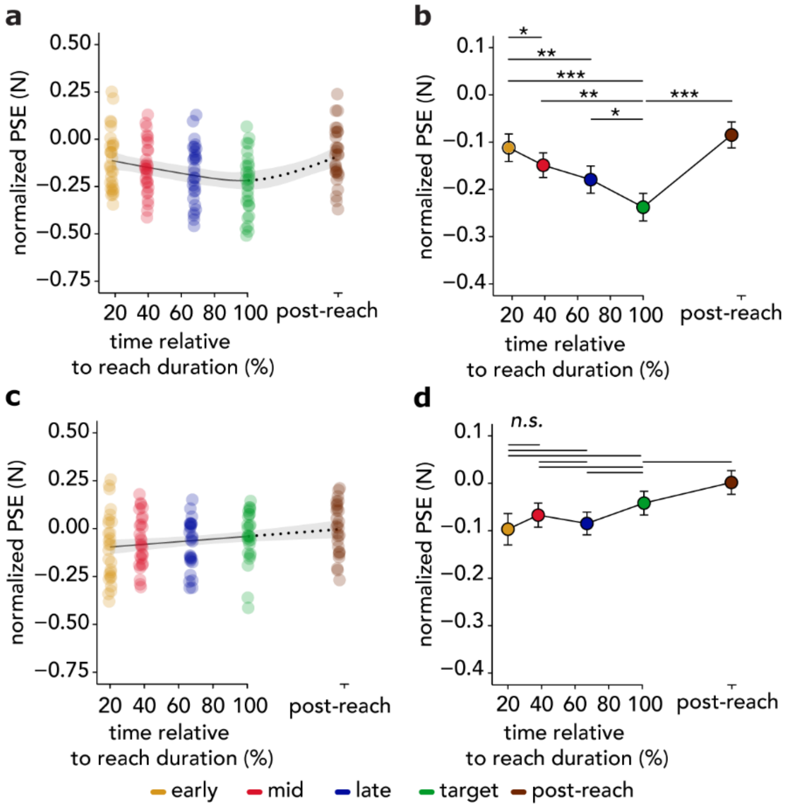


**Figure S7. Individual and group (mean ± s.e.m.) normalized PSEs for each type of trial in the *Self-touch* and *No self-touch* groups.** The *Self-touch* group **(a-b)** replicated all the effects observed in Experiment 1: touch felt progressively weaker as the reaching movement unfolded (*late* < *early*, *mid* < *early, target* < *early*, *target* < *mid*, *target* < *late*) and quickly recovered after the time of the predicted self-touch (*post-reach* > *target*). In contrast, the same comparisons yielded non-significant effects in the *No self-touch* group **(c-d)**. **(a, c)** Individual points are jittered (± 1%) to avoid complete overlapping. A regression line based on a generalized additive model is overlaid to illustrate the overall pattern in the data. The shaded area represents a 95% confidence interval (**Text S2**).

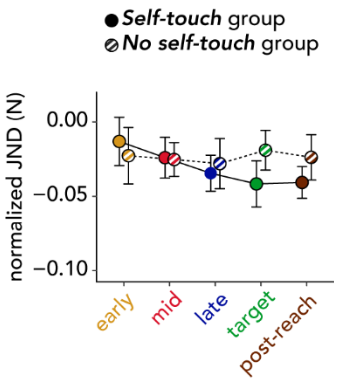


**Figure S8. Group (mean ± s.e.m.) normalized JNDs for each type of trial in the *Self-touch* group and *No self-touch* group.** There was no significant effect of trial type, indicating comparable sensory discrimination capacity between the two groups (**Text S3**).

### Text S1. Reaching movements in both experimental groups

The two groups performed reaching movements with similar duration (*Self-touch*: mean ± SD, 552 ± 61 ms, *No self-touch*: 571 ± 55 ms; $t(53.95) = -1.243$, $p = 0.219$, $CI^{95} = [-50.564, 11.860]$, $d = -0.332$; within-subjects standard deviation of movement duration, *Self-touch* group: 88 ± 20 ms, *No self-touch*: 72 ± 20 ms) (**Figure 3c**). The *No self-touch* group exhibited a slightly greater peak velocity (93 ± 11 cm/s) than the *Self-touch* group (85 ± 9 cm/s; $W = 580$, $p = 0.002$, $CI^{95} = [2.886, 13.631]$, $rrb = 0.481$) (**Figure 3d**), but both groups reached their peak velocities at comparable times (*Self-touch*: 217 ± 39 ms, *No Self-touch*: 206 ± 45 ms; $W = 449$, $p = 0.353$, $CI^{95} = [-10.702, 29.876]$, $rrb = 0.147$). The *test* forces were applied at similar times in the two groups: 99 ± 24 ms (18%), 213 ± 30 ms (39%), and 378 ± 31 ms (68%) in the *early*, *mid*, and *late* trials of the *Self-touch* group, and 116 ± 30 ms (20%), 219 ± 40 ms (38%), and 383 ± 24 ms (67%) in the *No Self-touch* group (**Figure 3e-f**). A mixed ANOVA on the times revealed, as expected, a significant main effect of trial type ($F(2, 108) = 5109.896$, $p < 0.001$, $\eta_p^2 = 0.990$), as well as a significant group-by-trial type interaction ($F(2, 108) = 3.177$, $p = 0.046$, $\eta_p^2 = 0.056$), but the post-hoc comparisons revealed no significant differences between the times when the *test* force was applied between the two groups (all *p*-values > 0.065 *FDR corrected).* There was no significant main effect of group ($F(1, 54) = 1.482$, $p = 0.229$, $\eta_p^2 = 0.027$). Finally, there were no significant differences in the forces applied by the participants in the *Self-touch* group (tap duration: 110 ± 39 ms) between the different reaching trial types, suggesting comparable taps between the trials ($F(1.77, 49.44) = 2.285$, p = 0.118, $\eta_p^2 = 0.075$) (**Figure S4**).

### Text S2. Complementary PSE analyses

In a complimentary analysis, we tested whether the findings of Experiment 1 were replicated by the *Self-touch* group that performed the same task. A repeated measures ANOVA on the normalized PSEs revealed a significant effect of trial type in the *Self-touch* group ($F(4, 112) = 13.633$, $p < 0.001$, $\eta_p^2 = 0.327$). Replicating the findings of Experiment 1, the PSEs were significantly weaker in the *mid* compared to *early* trials ($n = 29$, $t(28) = -2.172$, $p = 0.045$ *FDR-corrected*, $CI^{95} = [-0.072, -0.002]$, $d = -0.403$), in the *late* compared to the *early* trials ($n = 29$, $t(28) = -3.569$, $p = 0.003$ *FDR-corrected*, $CI^{95} = [-0.106, -0.029]$, $d = -0.663$), in the *target* compared to the *early* ($n = 29$, $t(28) = -4.927$, $p < 0.001$ *FDR-corrected*, $CI^{95} = [-0.178, -0.073]$, $d = -0.915$), in the *target* compared to the *mid* ($n = 29$, $t(28) = -3.479$, $p = 0.003$ *FDR-corrected*, $CI^{95} = [-0.141, -0.037]$, $d = -0.646$), and in the *target* compared to the *late* trials ($n = 29$, $t(28) = -2.750$, $p = 0.014$ *FDR-corrected*, $CI^{95} = [-0.102, -0.015]$, $d = -0.511$). As in Experiment 1, the difference in the PSEs between *mid* and *late* trials did not reach statistical significance ($n = 29$, $t(28) = 1.805$, $p = 0.082$ *FDR-corrected*, $CI^{95} = [-0.004, 0.065]$, $d = 0.335$). The lowest PSEs occurred at the time of contact between the two hands when the *test* force was perceived to be significantly weaker than *test* forces applied at all other probed times during the reaching. The PSEs quickly recovered in the *post-reach* trials and were significantly greater than in the *target* trials ($n = 29$, $t(28) = 5.704$, $p < 0.001$ *FDR-corrected*, $CI^{95} = [0.098,$

0.208], $d$ = 1.059). Together, the *Self-touch* group replicated all the effects observed in Experiment 1 with a new cohort and showed the same pattern of changes in somatosensory perception over time (**Figure S7a-b**).

We performed a similar analysis for the *No Self-touch* group. A main effect of the trial type was also revealed ($F$(2.76, 71.65) = 4.537, $p$ = 0.007, $\eta_p^2$ =0.149), but the PSEs did not change significantly as the reaching movement of the right hand progressed (*early* vs. *mid* trials: $n$ = 27, $t$(26) = −1.176, $p$ = 0.292 *FDR-corrected*, $CI^{95}$= [−0.081, 0.022], $d$ = −0.226; *early* vs *late* trials: $n$ = 27, $t$(26) = −0.409, $p$ = 0.686 *FDR-corrected*, $CI^{95}$= [−0.072, 0.048], $d$ = −0.079; *mid* vs. *late* trials: $n$ = 27, $t$(26) = 1.182, $p$ = 0.292 *FDR-corrected*, $CI^{95}$= [−0.013, 0.048], $d$ = 0.228; *late* vs. *target* trials: $n$ = 27, $t$(26) = −1.952, $p$ = 0.213 *FDR-corrected*, $CI^{95}$= [−0.088, 0.002], $d$ = −0.376). Moreover, the PSEs in the *target* trials were not significantly different from the PSEs in any other reaching trials (*target* vs. *early*: $n$ = 27, $t$(26) = 2.044, $p$ = 0.213 *FDR-corrected*, $CI^{95}$= [−3.219×10$^{-4}$, 0.110], $d$ = 0.393; *target* vs. *mid*: $n$ = 27, $t$(26) = 1.246, $p$ = 0.292 *FDR-corrected*, $CI^{95}$= [−0.017, 0.067], $d$ = 0.240) and did not significantly differ from the *post-reach* trials ($n$ = 27, $t$(26) = −1.753, $p$ = 0.213 *FDR-corrected*, $CI^{95}$= [−0.094, 0.007], $d$ = −0.337) (**Figure S7c-d**).To further provide evidence in favour of the absence of a decrease in the PSEs in the *No Self-touch* group, we conducted an additional Bayesian analysis in which we directly tested for a decrease in the PSEs during the reaching movement, as in the *Self-touch* group. The analysis consistently provided moderate to strong support for the absence of decreased PSEs between *early* and *mid* trials ($BF_{0+}$ = 9.766), *early* and *late* trials ($BF_{0+}$ = 6.512), *early* and *target* trials ($BF_{0+}$ = 13.394), *mid* and *target* trials ($BF_{0+}$ = 10.065) and *late* and *target* trials ($BF_{0+}$ = 13.028).

### Text S3. Control analyses

Given that the two groups differed in their peak velocity (**Text S1**), it is critical to test whether the observed PSE differences between the two groups were due to kinematic differences. Multiple linear regression analyses with the normalized PSEs as the dependent variable and peak velocity and group (*Self-touch* or *No self-touch*) as regressors revealed that the peak velocity was not a significant predictor of PSEs for *mid* ($b$ = 0.001, $p$ = 0.440), *late* ($b$ = 0.002, $p$ = 0.228), or *target* trials ($b$ = 0.001, $p$ = 0.599), thereby ruling out this alternative explanation.

We further tested whether the PSE differences were due to differences in task demands. For instance, concluding a reaching movement by tapping a sensor may be easier than stopping the movement precisely above a distance sensor. To test whether this was the case, we extracted the just noticeable differences (JND) for each participant for every trial type, normalized them to the *baseline* ($JND - JND_{baseline}$), and compared them between the two groups using a mixed ANOVA with the group as a between-subjects factor and type of trial as a within-subjects factor. The ANOVA revealed no significant main effect of trial type ($F$(4, 216) = 0.819, $p$ = 0.514, $\eta_p^2$ = 0.015), no significant main effect of group ($F$(1, 54) = 0.174, p = 0.678, $\eta_p^2$ = 0.003) and a non-significant group-by-trial type interaction ($F$(4, 216) = 0.970, $p$ = 0.425, $\eta_p^2$ = 0.018), showing that task difficulty was comparable between different trial types and between the *Self-touch* and *No self-touch* group (**Figure S8**).
